# Sequential delivery of LAIV and SARS-CoV-2 in the ferret model can reduce SARS-CoV-2 shedding and does not result in enhanced lung pathology

**DOI:** 10.1101/2021.02.01.429110

**Authors:** Kathryn A. Ryan, Katarzyna E. Schewe, Jonathan Crowe, Susan A. Fotheringham, Yper Hall, Richard Humphreys, Anthony C. Marriott, Jemma Paterson, Emma Rayner, Francisco J. Salguero, Robert J. Watson, Catherine J. Whittaker, Miles W. Carroll, Oliver Dibben

## Abstract

Co-circulation of SARS-CoV-2 and influenza viruses could pose unpredictable risks to health systems globally, with recent studies suggesting more severe disease outcomes in co-infected patients. The lack of a readily available COVID-19 vaccine has reinforced the importance of influenza vaccine programmes during the COVID-19 pandemic. Live Attenuated Influenza Vaccine (LAIV) is an important tool in protecting against influenza, particularly in children. However, it is unknown whether LAIV administration might influence the outcomes of acute SARS-CoV-2 infection or disease. To investigate this, quadrivalent LAIV (QLAIV) was administered to ferrets 3 days pre- or post-SARS-CoV-2 infection. LAIV administration did not exacerbate SARS-CoV-2 disease course or lung pathology with either regimen. Additionally, LAIV administered prior to SARS-CoV-2 infection significantly reduced SARS-CoV-2 replication and shedding in the upper respiratory tract (URT). We conclude that LAIV administration in close proximity to SARS-CoV-2 infection does not exacerbate mild disease and can reduce SARS-CoV-2 shedding.

## Introduction

Severe acute respiratory syndrome coronavirus 2 (SARS-CoV-2) emerged in late 2019, resulting in a pandemic and rapidly increasing the burden of respiratory illness globally. As of January 2021 the WHO reported over 98 million confirmed COVID-19 cases with more than 2 million deaths worldwide ^1^. The prospect of SARS-CoV-2 co-circulation with other respiratory viruses, particularly influenza, has raised major concerns over potential stresses to healthcare systems. A number of seasonal respiratory viruses including influenza A virus, influenza B virus and SARS-CoV-2 have been found in co-infected patients ^2–6^. SARS-CoV-2 and influenza viruses are easily transmissible within the population and the severity of disease caused in each case can range from asymptomatic to severe with a fatal outcome ^7,8^. However, the prevalence of COVID-19 and influenza co-infections rates varies across the literature ^9,10^ and the implications for disease severity are not fully understood. Stowe *et al.* retrospectively analyzed data from 19.256 UK patients during the first COVID-19 wave and showed that SARS-CoV-2 and influenza co-infections correlated with greater risk of severe illness and death, suggesting synergistic interaction between the viruses ^11^. Similar observations were made in the mouse model, where sequential influenza and SARS-CoV-2 infection resulted in exacerbated disease pathology ^12^.

Despite the recent approval of the first COVID-19 vaccines ^13–15^, pharmacological measures to control SARS-CoV-2 infection or COVID-19 disease remain limited. In the near-term, influenza vaccination remains of paramount importance in mitigating the risks associated with co-circulation of these viruses.

FluMist/Fluenz LAIV, is intranasally administered, with vaccine virus replication occurring in the epithelial cells of the upper respiratory tract, resulting in generation of both humoral and cellular immune responses ^16,17^. LAIV is primarily used in children, offering similar protection when compared with inactivated influenza vaccines ^18–23^. With the advantage of intranasal, needle free administration, LAIV has also demonstrated an ability to reduce community influenza transmission ^24^. LAIV can be administered to children with mild to moderate asthma as well as with mild fever, cold, runny nose or cough, as there is no evidence of LAIV exacerbating existing mild infections ^19,25–29^. However, the current COVID-19 pandemic gives rise to the possibility that LAIV could be administered to SARS-CoV-2 infected children displaying either mild or no COVID-19 symptoms ^30–32^. Alternatively, SARS-CoV-2 infection could be acquired following LAIV delivery. Due to the live, replication competent nature of LAIV, along with its intranasal delivery, it is feasible that co-incident LAIV and SARS-CoV-2 virus replication could take place. Currently there are no clinical data demonstrating this putative interaction with SARS-CoV-2 infection or the resulting COVID-19 illness in children.

The ferret has been shown to provide a model of mild COVID-19 disease with consistent upper respiratory tract shedding of SARS-CoV-2 ^33–35^, thus offering a surrogate for the mild COVID-19 disease seen in pediatric infections. Using the ferret model we aimed, firstly, to describe the impact of LAIV administration on pre-existing, acute SARS-CoV-2 infection and associated respiratory pathology. Secondly, we have evaluated the effect of LAIV given prior to SARS-CoV-2 infection. Overall we intended to understand whether, in either scenario, LAIV might exacerbate or ameliorate SARS-CoV-2 infection and disease.

## Results

To assess the implications of LAIV and SARS-CoV-2 co-infection and its impact on COVID-19 disease severity, SARS-CoV-2 at 6.7 log_10_ PFU per dose and quadrivalent LAIV (QLAIV) at 4.0 log_10_ FFU per dose, per strain were administered intranasally to groups of six ferrets (Figure 1a). Optimization of these SARS-CoV-2 and QLAIV doses was described by Ryan *et al.* ^35^ and Dibben *et al.* ^36^, respectively. LAIV administration occurred either 3 days before (LAIV/SARS-CoV-2) or 3 days after (SARS-CoV-2/LAIV) the SARS-CoV-2 challenge (Figure 1b). Control groups for SARS-CoV-2 only (Mock/SARS-CoV-2) and LAIV only (LAIV/Mock) were also included. All study groups are summarized in Figure 1c. The endpoints of the study were the histopathological examination of lung and nasal cavity tissues and the quantification of SARS-CoV-2 virus RNA in throat swabs, nasal cavity tissues and lung tissues.

**Figure 1.**
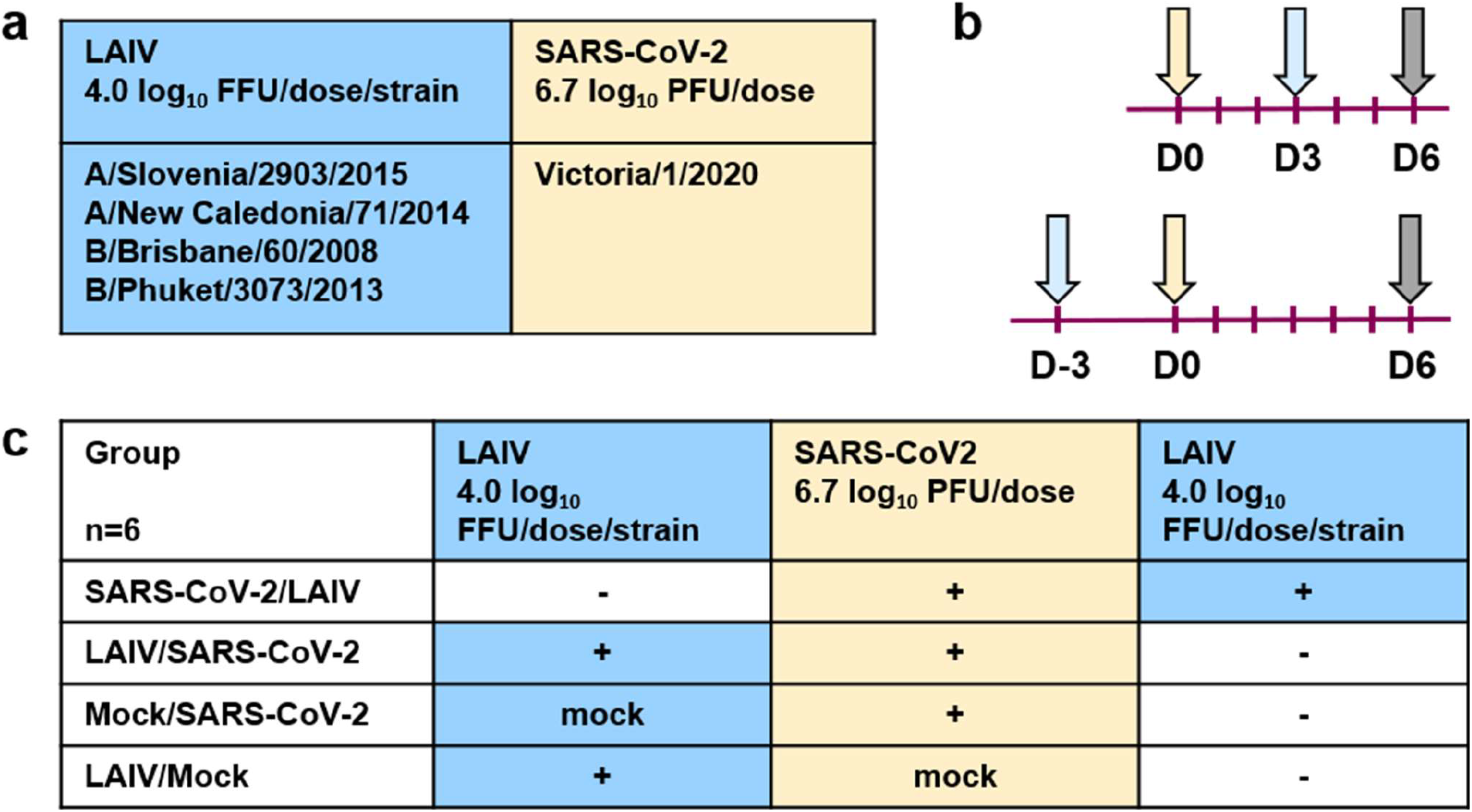
Study design for assessment of impact of LAIV on SARS-CoV-2 infection in ferrets. (a) Quadrivalent LAIV formulation with 4.0 log_10_ FFU/dose/strain and SARS-CoV-2 with 6.7 log_10_ PFU/dose were utilized here. (b) LAIV and SARS-CoV-2 were sequentially administered to ferrets intranasally. Yellow arrows indicate the SARS-CoV-2 infection at day 0 (d0) and light blue arrows indicate LAIV administration (d3 or d-3). Vertical burgundy lines indicate throat swabs at days 1, 2, 3, 4, 5 and 6. Grey arrows indicate cull and tissue harvest. (c) Four groups of six ferrets were included. LAIV was administered either post SARS-CoV-2 challenge (SARS-CoV-2/LAIV) or pre SARS-CoV-2 challenge (LAIV/SARS-CoV-2). Control groups received either mock LAIV and SARS-CoV-2 (Mock/SARS-CoV-2) or LAIV and mock SARS-CoV-2 (LAIV/Mock).

### Histopathological examination shows no effect of LAIV on SARS-CoV-2 lung pathogenesis and only a mild increase in the nasal cavity

All ferrets were euthanized 6 days after SARS-CoV-2 challenge. Pathological changes were assessed in all lung lobes and the nasal cavity.

In the nasal cavity tissues, inflammatory and degenerative changes were observed in the mucosa which were similar in appearance for all animals in all groups (Figure 2). These comprised mixed inflammatory cells, mainly lymphocytes, infiltrating the propria-submucosa and neutrophils infiltrating the epithelial layer, with variable presence and severity of epithelial degeneration, loss and areas of denudation. Lesions were present in both the rostral and caudal areas of the nasal cavity, containing both the respiratory and olfactory epithelia, without a detectable predilection for a specific location. Patchy exudates, comprising degenerate neutrophils, epithelial cells and mucus, were also noted. These inflammatory and degenerative changes were most prominent in animals that had either received LAIV post-SARS-CoV-2 challenge (SARS-CoV-2/LAIV) (Figure 2a) or LAIV pre-SARS-CoV-2 challenge (LAIV/SARS-CoV-2) (Figure 2b). Severity was slightly reduced in animals that had either received a mock vaccine prior to SARS-CoV-2 challenge (Mock/SARS-CoV-2) (Figure 2c) or LAIV prior to a mock challenge (LAIV/Mock) (Figure 2d).

**Figure 2.**
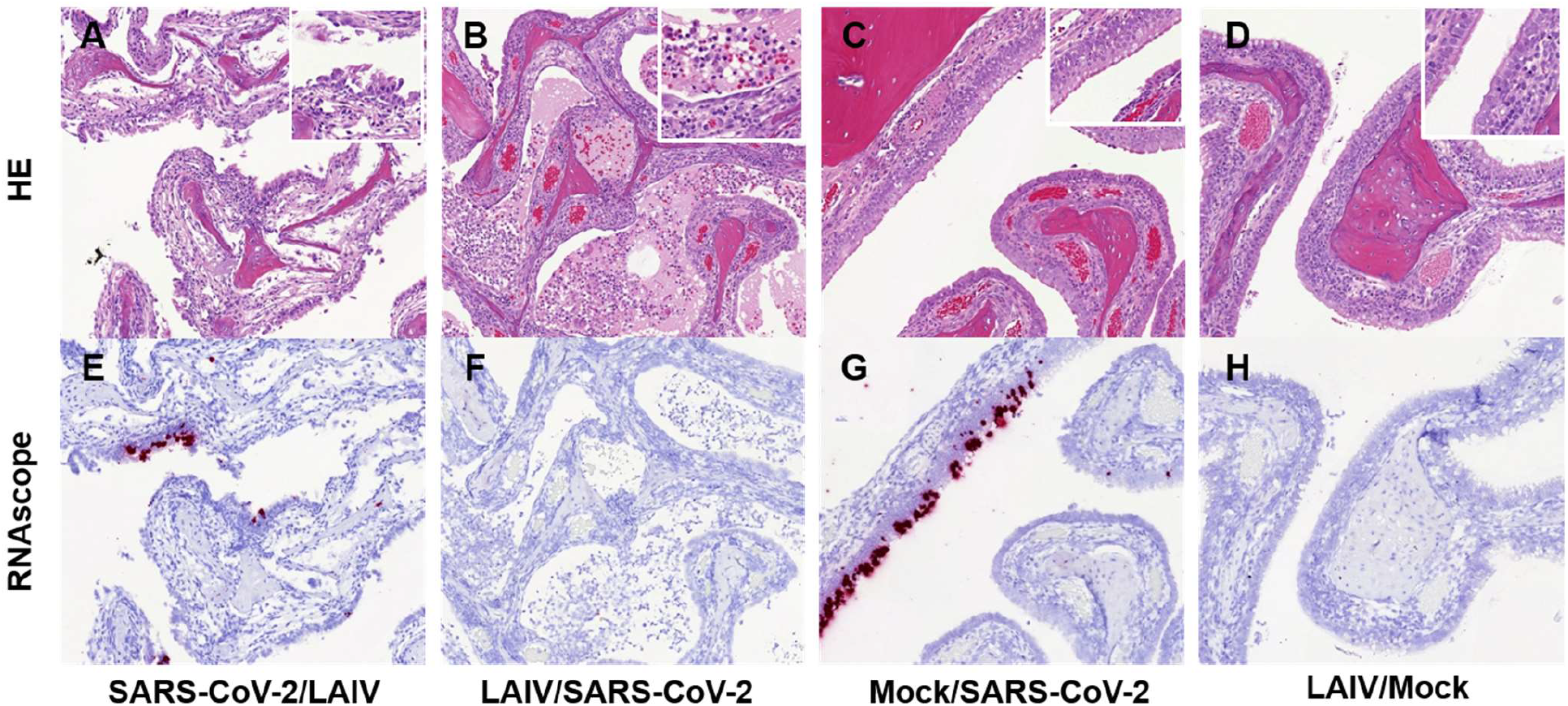
Histopathological changes and SARS-CoV-2 RNA detection in the nasal cavity. A) Group SARS-CoV-2/LAIV; B) Group LAIV/SARS-CoV-2; C) Group Mock/SARS-CoV-2 and D) Group LAIV/Mock. Microscopic inflammatory changes with variable epithelial degeneration and loss and exudate. 200 × magnification, insets 823 × magnification. (HE). Viral staining for SARS-CoV-2 RNA is positive in E) Group SARS-CoV-2/LAIV; and G) Group Mock/SARS-CoV-2, and absent in F) Group LAIV/SARS-CoV-2; and H) Group LAIV/Mock. 200 × magnification.

Variable levels of positive staining for the presence of SARS-CoV-2 RNA were detected in epithelial and sustentacular cells, and in cellular exudates in five of the six animals that had received LAIV post-SARS-CoV-2 challenge (SARS-CoV-2/LAIV) (Figure 2e) and in four out of six animals that had received mock vaccine pre-SARS-CoV-2 challenge (Mock/SARS-CoV-2) (Figure 2g). Staining was absent in all animals that received LAIV pre-SARS-CoV-2 challenge (LAIV/SARS-CoV-2) (Figure 2f) and in all LAIV only control animals (LAIV/Mock) (Figure 2h).

In the lungs, microscopic inflammatory changes involving both parenchyma and airways were observed in all animals from all groups (Supplementary Figure 1); all lobes on both left and right sides were affected with a multifocal patchy to diffuse distribution of lesions. Within the parenchyma there were scattered foci of inflammatory cells comprising primarily of mononuclear cells with occasional neutrophils, filling alveolar spaces and expanding walls (multi-focal alveolitis). Lymphocytes were also observed surrounding blood vessels (perivascular lymphocytic cuffing). In both the bronchi and bronchioles, a mixed inflammatory cell infiltration was seen, with neutrophils and occasionally eosinophils infiltrating the epithelium, and mononuclear cells accumulating both within the walls and in peribronchiolar locations (bronchitis and bronchiolitis). Degenerate polymorphonuclear leukocytes, cellular debris and mucus were noted in the lumen of some airways.

The overall severity of these lesions was low to mid-grade and similar between all four groups (Figure 3a and 3b), with perivascular lymphocytic cuffing and bronchiolitis representing the most prominent changes. No significant differences were observed between the lung pathology scores of any of the groups (Figure 3b). The nasal cavity tissue pathology scores were summed to enable statistical analysis (Figure 3c). No significant difference was observed between the Mock/SARS-CoV-2 and LAIV/Mock groups. The SARS-CoV-2/LAIV and LAIV/SARS-CoV-2 group scores were significantly higher than both the Mock/SARS-CoV-2 group (p=0.0443 and p=0.0012, respectively) and the LAIV/Mock group (p=0.0030 and p<0.0001, respectively). However, their severity was still classified as mild to moderate.

**Figure 3.**
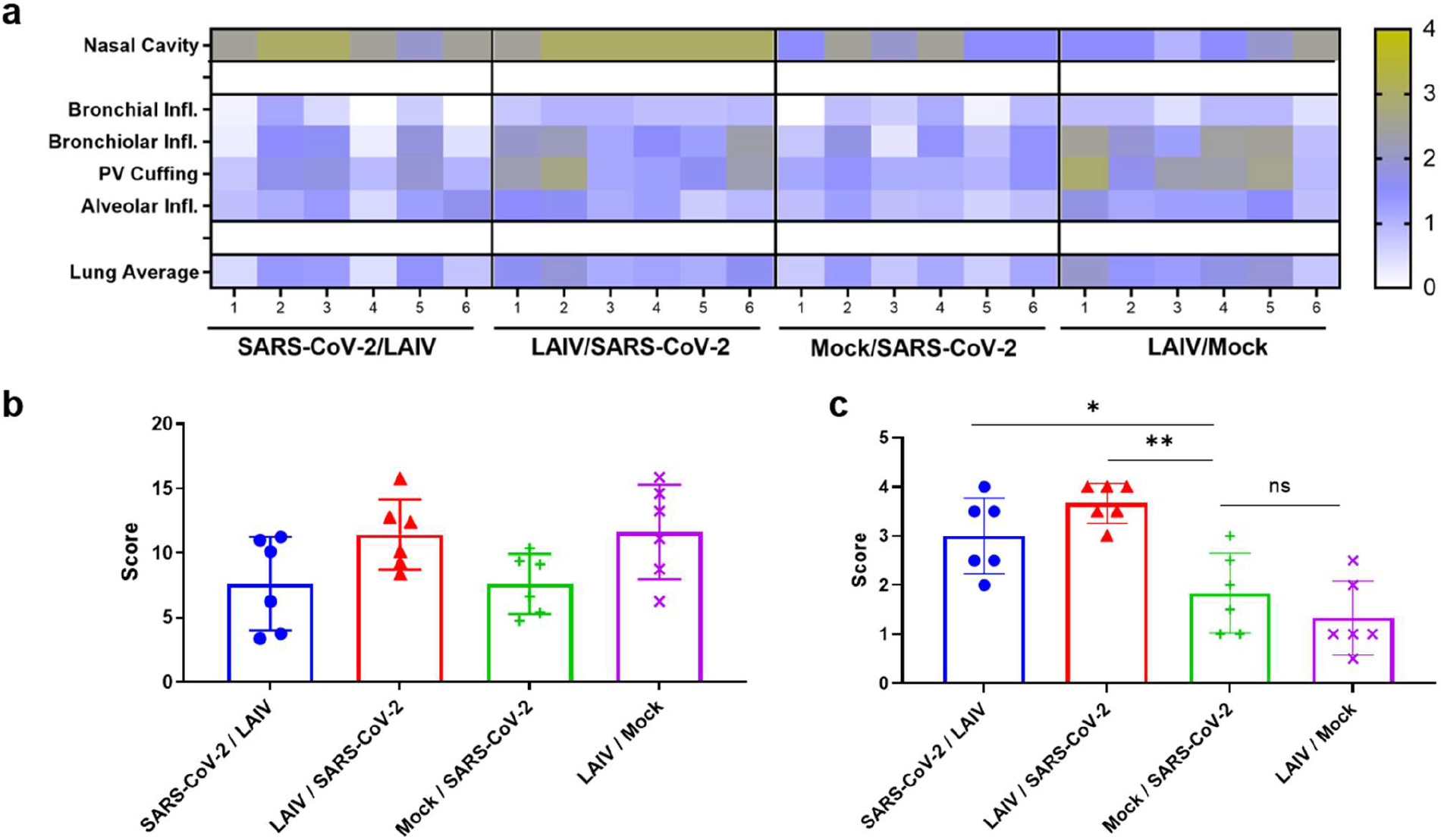
Histopathological examination of lung and nasal cavity tissue shows no impact of LAIV on SARS-CoV-2 lung pathology. Lung and nasal cavity tissue samples were taken from all ferrets at day 6 post challenge and inflammation related parameters were examined: epithelial inflammation and degeneration in nasal cavity tissue and inflammation (Infl.) of alveolar walls/spaces, perivascular (PV) lymphocytic cuffing, bronchiolar inflammation and bronchial inflammation in lung tissue. (a) Changes were scored from 0 (normal), 1 (minimal), 2 (mild), 3 (moderate) to 4 (marked) for each ferret and summarized in the heat map. (b) Lung scores were compared. Symbols indicate the sum of pathology scores per ferret and bars show the group means with standard deviation. No significant differences were noted between any groups. (c) Nasal cavity tissue scores were compared. Symbols indicate epithelial inflammation and degeneration in nasal cavity tissue and bars show the group means with standard deviation. No significant difference was observed between both control groups (Mock/SARS-CoV-2 and LAIV/Mock). SARS-CoV-2/LAIV and LAIV/SARS-CoV-2 scores were significantly higher than Mock/SARS-CoV-2 (p=0.0443 and p=0.0012, respectively) and higher than LAIV/Mock (p=0.0030 and p<0.0001, respectively). Statistical significance is shown by horizontal lines and P-value indicators: ^ns^P > 0.05, *P < 0.05, **P < 0.01.

SARS-CoV-2 RNA was not detected in the lungs of any animal in any group. Summaries of the presence and severity of viral RNA in the nasal cavity tissue and severity scores for microscopic lesions in the nasal cavity tissue are shown in Figure 2, Figure 3 and Supplementary Table 1.

These observations showed that co-administration of LAIV and SARS-CoV-2 did not result in enhanced lung pathogenesis and that SARS-CoV-2 RNA levels in co-infected ferrets were not enhanced when compared to SARS-CoV-2 only controls (Mock/SARS-CoV-2).

### Viral shedding post challenge in throat swabs is reduced in animals receiving LAIV before SARS-CoV-2

The viral RNA of SARS-CoV-2 was detected in throat swabs from day 1 post challenge (pc) in SARS-CoV-2 only treated ferrets (Mock/SARS-CoV-2). Shedding reached a peak at day 3 pc, with a titre of 5.85 log_10_ copies/ml and remained consistent until day 6 pc (Supplementary Figure 2). To enable statistical comparison between groups, shedding for individual animals was expressed as the geometric mean of shedding per day, with group means shown in Figure 4a. For the Mock/SARS-CoV-2 group mean shedding was 4.98 log_10_ copies/ml/day. The mean shedding in LAIV primed ferrets (LAIV/SARS-CoV-2) was significantly reduced, by a factor of 7, to 4.13 log_10_ copies/ml/day (p=0.0059), while no significant change was noted in SARS-CoV-2/LAIV ferrets (4.65 log_10_ copies/ml/day (p=0.51) (Figure 4a).

**Figure 4.**
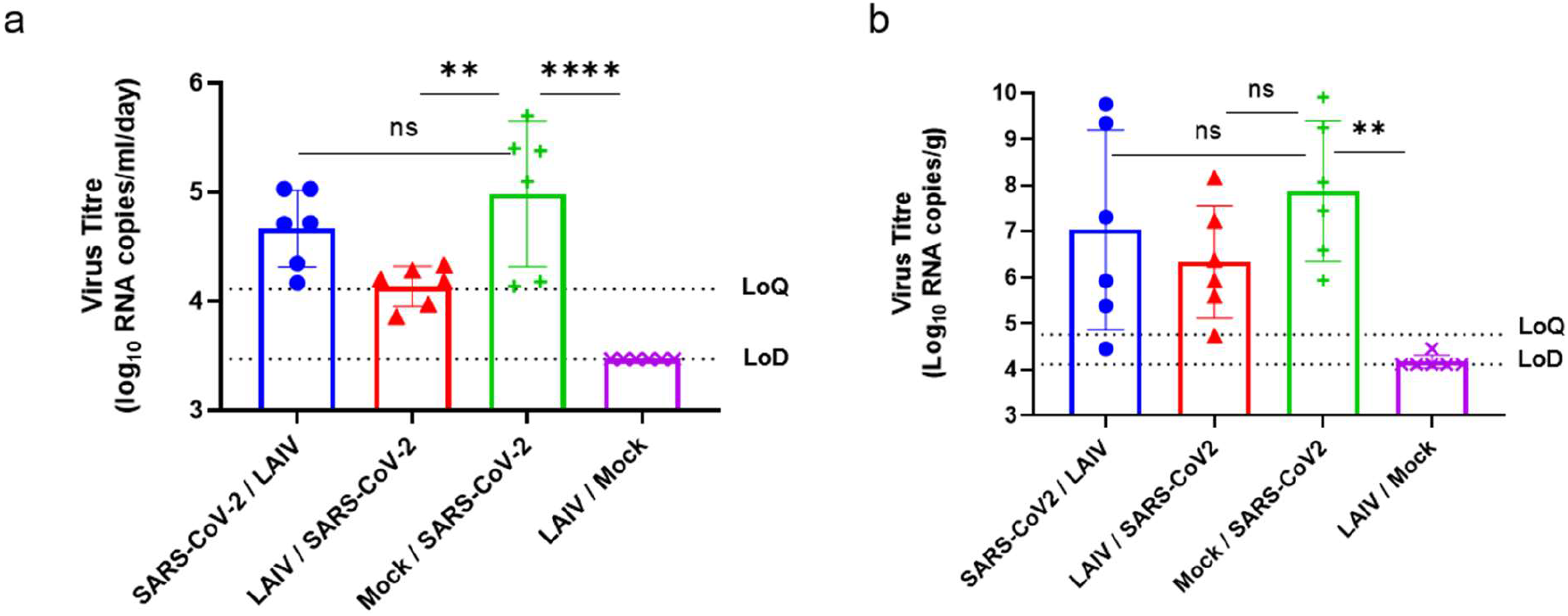
SARS-CoV-2 viral RNA detection in throat swabs and nasal cavity tissue. (a) The throat swabs were taken at days 1, 2, 3, 4, 5 and 6 pc and SARS-CoV-2 viral loads were determined by RT-qPCR. The symbols show the geometric mean of virus titres over 6 days for one ferret, bars represent the group mean of 6 ferret with standard deviation. Statistical significance are shown by horizontal lines and P-value indicators: ^ns^P > 0.05, **P < 0.01, ****P < 0.0001. Horizontal dotted lines show the lower limit of quantification (LoQ=4.11 log_10_ RNA copies/ml/day) and the lower limit of detection (LoD=3.47 log_10_ RNA copies/ml/day). SARS-CoV-2 shedding in group Mock/SARS-CoV-2 was significantly higher than in group LAIV/SARS-CoV-2 (p=0.0059) and in group LAIV/Mock (p<0.0001). No significant changes were noted between group SARS-CoV-2/LAIV and control group Mock/SARS-CoV-2 (p=0.51). (b) The SARS-CoV-2 RNA was measured in nasal cavity tissue homogenates by RT-qPCR. The symbols show the virus titre measured in nasal cavity tissue at day 6 for one ferret, bars represent the group mean of 6 ferret with standard deviation. Statistical significance is shown by horizontal lines and P-value indicators: ^ns^P > 0.05, **P < 0.01. Horizontal dotted lines show the lower limit of quantification (LoQ=4.76 log_10_ RNA copies/g) and the lower limit of detection (LoD=4.12 log_10_ RNA copies/g). SARS-CoV-2 shedding in group Mock/SARS-CoV-2 was significantly higher than in group LAIV/Mock (p=0.0015). No statistical differences between SARS-CoV-2 only control group (Mock/SARS-CoV-2) and both co-infected groups LAIV/SARS-CoV-2 and SARS-CoV-2/LAIV were noted in nasal cavity tissue (p=0.2924 and p=0.7552, respectively).

### Viral loads in tissues

Viral RNA from SARS-CoV-2 was detected in lung and nasal cavity tissue homogenates. Quantifiable levels of SARS-CoV-2 viral RNA were not detected in the lungs of any ferret at 6 days post-SARS-CoV-2 challenge. SARS-CoV-2 viral RNA was detected in the nasal cavity tissue of all ferrets challenged with SARS-CoV-2. SARS-CoV-2/LAIV animals showed no difference in nasal cavity RNA levels relative to the Mock/SARS-CoV-2 group, while the LAIV/SARS-CoV-2 group showed a trend for lower levels of viral RNA in the nasal cavity tissue when compared to both SARS-CoV-2/LAIV and Mock/SARS-CoV-2 groups. However, this difference was not statistically significant. Ferrets in the LAIV/Mock group were confirmed negative for SARS-CoV-2 viral RNA (Figure 4b).

Taken together, these viral RNA data showed no significant changes in SARS-CoV-2 virus replication if LAIV was administered 3 days post-SARS-CoV-2 challenge (SARS-CoV-2/LAIV) relative to the Mock/SARS-CoV-2 control group. LAIV administration 3 days pre-SARS-CoV-2 challenge resulted in a reduction of SARS-CoV-2 virus replication in the LAIV/SARS-CoV-2 group when compared to SARS-CoV-2 only (Mock/SARS-CoV-2).

## Discussion

In order to investigate the effect of LAIV on SARS-CoV-2 replication and pathology we applied the ferret model for mild SARS-CoV-2 clinical disease described by Ryan *et al.* ^35^. In SARS-CoV-2 infected control animals, mild histopathological changes were observed in respiratory tract tissues, corresponding to the mild disease pattern described previously ^35^. For all animals in the SARS-CoV-2 only group (Mock/SARS-CoV-2), microscopic lesions were present with greatest prominence in the nasal cavity tissue and comprised of acute to sub-acute, inflammatory and degenerative changes. Histopathological changes were also seen in the lungs, but with a lower frequency and severity. These findings agreed with previous challenge study data and reflected a reduced severity of pathological changes in the lower respiratory tract (LRT) compared to the URT ^35^. Additionally, the mild disease phenotype observed in ferrets appeared to model the mild COVID-19 progression in children ^30–32^.

Mild histopathological changes in both the lungs and nasal turbinates were also noted in animals that received LAIV only (LAIV/Mock). LAIV has an excellent safety profile in clinical studies ^19,29^, suggesting that this level of pathology resulted from LAIV induced immune response.

Microscopic lesions in lungs of co-infected ferrets (LAIV/SARS-CoV-2 and SARS-CoV-2/LAIV) were primarily low-grade and similar in appearance to the control groups, irrespective of the infection schedule, suggesting that LAIV did not alter the LRT disease phenotype if administered either 3 days pre- or post-SARS-CoV-2 infection. Co-infected ferrets did show significantly higher pathology scores in nasal cavity tissue relative to both the SARS-CoV-2 only and LAIV only control groups. This likely reflected the combined immune response to SARS-CoV-2 challenge and LAIV administration. Despite this, pathology in the URT remained mild and, coupled with the absence of observable changes in the lungs, was not considered indicative of an enhanced disease phenotype.

SARS-CoV-2 RNA was not detected by RT-qPCR in the lungs of any animal in any group, these findings agree with the mild pathology observed. In the SARS-CoV-2 control animals (Mock/SARS-CoV-2), SARS-CoV-2 RNA was readily detected by RT-qPCR in throat swabs and nasal cavity tissues, confirming previous observations ^35^. SARS-CoV-2 RNA was also detected in throat swabs and nasal turbinate tissues of co-infected animals (SARS-CoV-2/LAIV and LAIV/SARS-CoV-2). However, while RNA levels in the SARS-CoV-2/LAIV group were similar to the SARS-CoV-2 only group (Mock/SARS-CoV-2), SARS-CoV-2 shedding was significantly reduced in animals that received LAIV prior to SARS-CoV-2 challenge (LAIV/SARS-CoV-2). These observations aligned with RNAscope observations of nasal cavity tissue, where RNA staining was present in the SARS-CoV-2 only (Mock/SARS-CoV-2) and SARS-CoV-2/LAIV groups but not in the LAIV/SARS-CoV-2 or LAIV only (LAIV/Mock) groups. Although no significant differences were seen between SARS-CoV-2 RNA levels in nasal cavity tissue homogenates by RT-qPCR, a trend for a reduction in RNA levels in LAIV/SARS-CoV-2 animals was also evident.

Taken together, these data suggested that the administration of LAIV did not enhance SARS-CoV-2 RNA shedding in ferrets. Moreover, administration of LAIV 3 days prior to SARS-CoV-2 challenge resulted in a reduction in SARS-CoV-2 replication and shedding in the ferret URT. A reduction in SARS-CoV-2 shedding in the clinic could have important implications for transmission across the population ^37^.

Our findings of reduced SARS-CoV-2 replication and shedding in animals receiving LAIV prior to SARS-CoV-2 align with recent observations in the mouse model, where SARS-CoV-2 viral load in the lungs was significantly reduced following an initial wild type influenza virus infection, when compared to SARS-CoV-2 only infected animals ^12^. However, influenza/SARS-CoV-2 co-infected mice displayed more severe disease progression despite reduced SARS-CoV-2 viral load ^12^. This is contrary to our observations for LAIV, showing no evidence of enhancement of mild lung pathology in LAIV/SARS-CoV-2 co-infected ferrets. This was likely due to the higher pathogenicity of wild type influenza virus relative to LAIV, in concert with a more severe COVID-19 disease model in mice.

We hypothesize that viral competition might play a role in the interaction of LAIV and SARS-CoV-2, as well as LAIV-mediated non-specific immune response. Previously, LAIV has been shown to provide non-specific immediate protection against Respiratory Syncytial Virus infection in mice ^38^. In another mouse model LAIV achieved immediate, broad-spectrum protection against heterologous influenza strains in the absence of specific antibodies, mediated by antiviral IFN-α responses and pro-inflammatory cytokines ^39^. Similar observations were made in the clinic, where LAIV protected children against an influenza strain that was antigenically distinct from the vaccine strain, within two weeks of vaccination, even in children who only received their initial priming dose of LAIV ^40^. Zhu *et al.* investigated transcriptomic changes in whole blood of LAIV vaccinated children; at seven days post vaccination they identified clusters of co-expressed genes either modulated by type 1 interferons (IFN) or involved in IFN-regulated pathways. These findings support the establishment of LAIV-induced, IFN-mediated non-specific antiviral immunity in the first few weeks post vaccination ^41^. The concept of administering an unrelated live vaccine, such as MMR, as a preventive measure against COVID-19 has been proposed for similar reasons ^42,43^. LAIV-induced, non-specific immunity as a means of passive protection against SARS-CoV-2 infection could therefore prove feasible. However, the amplitude and duration of any LAIV mediated protection is not understood and more clinical data on COVID-19 disease in the context of LAIV uptake could further this understanding.

Whether driven by direct viral competition or activation of non-specific immune responses, LAIV dependent reduction in SARS-CoV-2 shedding is likely to be dependent on the proximity of the two infections. Here, a 3 day interval was used with the aim of achieving peak replication of the first infection by the time of administration of the second. With longer intervals between infections it is possible that this inhibitory effect would decline, due to either the waning of non-specific immunity or to a loss of viral competition. Further work is required to understand the longevity of this inhibition.

At the same time the possible impact of SARS-CoV-2 infection on LAIV effectiveness must be considered. While this was outside the scope of the work described here, investigations into this important question are ongoing.

Our findings deliver the first evidence that LAIV does not exacerbate the mild pathogenic changes caused by SARS-CoV-2 in the LRT of ferrets when delivered shortly pre- or post-challenge. Moreover, if administered 3 days pre-SARS-CoV-2, LAIV reduced SARS-CoV-2 viral replication and shedding in the URT, raising the possibility of a reduction in transmission as a result.

This work supports the administration of LAIV to children of unknown COVID-19 status and suggests a potential additional benefit of LAIV administration during the COVID-19 pandemic.

## Methods

### Cells and viruses

SARS-CoV-2 strain Victoria/01/2020 ^44^ was generously provided by The Doherty Institute, Melbourne, Australia at P1 after primary growth in Vero/hSLAM cells and subsequently passaged twice at Public Health England (PHE) Porton in Vero/hSLAM cells (ECACC 04091501). Cells were infected with ~0.0005 MOI of virus and harvested at day 4 by a single freeze thaw cycle followed by clarification by centrifugation at 1000 × g for 10 min. Whole genome sequencing was performed on the P3 challenge stock using both Nanopore and Illumina as described previously ^45^. Virus titre of the challenge stocks was determined by plaque assay on Vero/E6 cells (ECACC 85020206). Cell lines were obtained from the European Collection of Authenticated Cell Cultures (ECACC) PHE, Porton Down, UK. Cell cultures were maintained at 37 °C in MEM (Life Technologies, California, USA) supplemented with 10% fetal bovine serum (Sigma, Dorset, UK) and 25 mM HEPES (Life Technologies). In addition, Vero/hSLAM cultures were supplemented with 0.4 mg/ml of geneticin (Invitrogen) to maintain the expression plasmid.

The quadrivalent LAIV formulation used was representative of the 2017-18 vaccine composition and was generated from non-commercial material for use in these studies. LAIV viruses were propagated in the allantoic cavity of 10–11-day-old embryonated hens’ eggs (Charles River Laboratories, Wilmington, MA, USA) at 33 °C. All were 6:2 reassortants carrying the six internal gene segments (PB2, PB1, PA, NP, M, and NS) of cold-adapted A/AnnArbor/6/1960 ^46^ or cold-adapted B/AnnArbor/1/1966 ^47^ and the HA and neuraminidase (NA) gene segments of the following viruses: A/H1N1pdm09 – A/Slovenia/2903/2015, H3N2 – A/New Caledonia/71/2014, B/Yamagata – B/Phuket/3073/2013, B/Victoria – B/Brisbane/60/2008. LAIV was titrated by focus-forming assay (FFA) as previously described ^48^. Madin–Darby canine kidney (MDCK) used in FFA, were obtained from American Type Culture Collection (ATCC) and passaged fewer than 20 times prior to use in analytical tests. MDCK cells were cultured and maintained in Eagle’s minimum essential medium with non-essential amino acids at 37 °C and 5% CO_2_, as previously described ^49^.

### Ferrets

The work described here was conducted at Porton Down, Public Health England (PHE). Twenty-four healthy, female ferrets (*Mustela putorius furo*) aged 7 months were obtained from a UK Home Office accredited supplier (Highgate Farm, UK). The mean weight at the time of challenge was 1006g/ferret (range 829-1225g). Ferrets were housed at Advisory Committee on Dangerous Pathogens (ACDP) containment level 2 in social groups of 2-6 or at ACDP containment level 3 animals were housed in pairs. Cages met with the UK Home Office *Code of Practice for the Housing and Care of Animals Bred, Supplied or Used for Scientific Procedures* (December 2014). Access to food and water was *ad libitum* and environmental enrichment was provided. All experimental work was conducted under the authority of a UK Home Office approved project license that had been subject to local ethical review at PHE Porton Down by the Animal Welfare and Ethical Review Body (AWERB) as required by the *Home Office Animals (Scientific Procedures) Act 1986*.

Ferrets were confirmed seronegative for circulating H1N1, H3N2, and B viruses by hemagglutination inhibition (HAI) assay and randomly assigned to study groups.

### Study Design

On Study Day 0 the ferrets were lightly sedated with isoflurane and intranasally challenged with a single 1.0 ml (~0.5 ml/naris) dose of SARS-CoV-2. Groups received either mock challenge (phosphate buffered saline, Gibco, cat. 10010023) or SARS-CoV-2 at dose of 6.7 log_10_ PFU/animal in the same sample diluent. LAIV was administered intranasally at day −3 or 3 with a single 0.2 ml (~0.1 ml/naris) dose of LAIV. Groups received either mock vaccination (phosphate buffered saline (ThermoFisher Scientific, Waltham, Massachusetts, USA) with 1x sucrose phosphate (ThermoFisher Scientific: custom product, cat. AC10210390) and 1x gelatine-arginine-glutamate (ThermoFisher Scientific: custom product, cat. AC10207676) or LAIV at a dose of 4.0 log_10_ FFU/strain/animal, in the same sample diluent.

To evaluate SARS-CoV-2 virus shedding following challenge, throat swabs were taken daily on days one to six post SARS-CoV-2 challenge. Animals were sedated with isoflurane and a flocked swab (Copan, cat. 552C) was gently stroked across the back of the pharynx in the tonsillar area. Swabs were eluted by light vortexing in 1 ml of Copan universal transport medium (UTM) (Copan, cat. 330C), aliquoted and stored at –80 °C.

Ferrets from all groups were euthanized at 6 days post SARS-CoV-2 challenge. Ferrets were anaesthetized with an intramuscular injection of ketamine/xylazine (17.9 mg/kg and 3.6 mg/kg bodyweight) and exsanguination was effected via cardiac puncture, followed by injection of an anesthetic overdose (sodium pentabarbitone Dolethal, Vetquinol UK Ltd, 140 mg/kg). A necropsy was performed immediately after confirmation of death. Lower left lung lobe and nasal turbinate were collected into RNAprotect (Qiagen, cat. 76163). Remaining lung tissue and nasal cavity tissues were placed in 10% neutral buffered formalin for histopathological analysis.

### Pathological Studies

Samples of lung lobe and nasal cavity tissue were fixed by immersion in 10% neutral-buffered formalin, processed and embedded into paraffin wax. Nasal cavity samples were decalcified using an EDTA-based solution prior to embedding. Sections of 4 μm were cut and stained with hematoxylin and eosin (HE) and examined microscopically. A nasal cavity and lung tissue histopathology scoring system was used to evaluate the severity of histopathological lesions observed in each animal (Supplementary Table 2). Sections were analyzed by two qualified veterinary pathologists. In addition, samples were stained using the RNAscope technique to identify the SARS-CoV-2 virus RNA. Briefly, tissues were pre-treated with hydrogen peroxide for 10 minutes (room temperature), target retrieval for 15 mins (98-101 °C) and protease plus for 30 mins (40 °C) (Advanced Cell Diagnostics). A V-nCoV2019-S probe (Advanced Cell Diagnostics, cat. 848561) was incubated on the tissues for 2 hours at 40 °C. Amplification of the signal was carried out following the RNAscope protocol using the RNAscope 2.5 HD Detection kit – Red (Advanced Cell Diagnostics). Digital image analysis was performed on RNAscope labelled slides to ascertain the percentage of stained cells within the lesions, using the Nikon-NIS-Ar package. One section per lung lobe and 2 nasal cavity tissue sections (including rostral and caudal areas) were assessed. Nasal cavity tissue sections covered the whole structure (transversal section, both left and right).

### RT-qPCR

#### RNA extraction

RNA was isolated from nasal swabs and throat swabs. Samples were inactivated in AVL (Qiagen) and ethanol. Downstream extraction was then performed using the BioSprint™96 One-For-All vet kit (Indical) and Kingfisher Flex platform as per manufacturer’s instructions. Tissues were homogenized in Buffer RLT+ beta-mercaptoethanol (Qiagen). Tissue homogenate was then centrifuged through a QIAshredder homogenizer (Qiagen) and supplemented with ethanol as per manufacturer’s instructions. Downstream extraction from tissue samples was then performed using the BioSprint™96 One-For-All vet kit (Indical) and Kingfisher Flex platform as per manufacturer’s instructions.

#### SARS-CoV-2 Viral RNA Quantification

Reverse transcription-quantitative polymerase chain reaction (RT-qPCR) targeting a region of the SARS-CoV-2 nucleocapsid (N) gene was used to determine viral loads and was performed using TaqPath™ 1-Step RT-qPCR Master Mix, CG (Applied Biosystems™), 2019-nCoV CDC RUO Kit (Integrated DNA Technologies) and QuantStudio™ 7 Flex Real-Time PCR System. Sequences of the N1 primers and probe were: 2019-nCoV_N1-forward, 5’ GACCCCAAAATCAGCGAAAT 3’; 2019-nCoV_N1-reverse, 5’ TCTGGTTACTGCCAGTTGAATCTG 3’; 2019-nCoV_N1-probe, 5’ FAM-ACCCCGCATTACGTTTGGTGGACC-BHQ1 3’. The cycling conditions were: 25 °C for 2 minutes, 50 °C for 15 minutes, 95 °C for 2 minutes, followed by 45 cycles of 95 °C for 3 seconds, 55 °C for 30 seconds. The quantification standard was *in vitro* transcribed RNA of the SARS-CoV-2 N ORF (accession number NC_045512.2) with quantification between 1 and 6 log_10_ copies/μl. Positive swab and fluid samples detected below the limit of quantification (LoQ) of 4.11 log_10_ copies/ml, were assigned the value of 5 copies/μl, this equates to 3.81 log_10_ copies/ml, whilst undetected samples were assigned the value of < 2.3 copies/μl, equivalent to the assay’s lower limit of detection (LoD) which equates to 3.47 log_10_ copies/ml. Positive tissue samples detected below the limit of quantification (LoQ) of 4.76 log_10_ copies/ml were assigned the value of 5 copies/μl, this equates to 4.46 log_10_ copies/g, whilst undetected samples were assigned the value of < 2.3 copies/μl, equivalent to the assay’s lower limit of detection (LoD) which equates to 4.76 log_10_ copies/g.

### Quantification and statistical analysis

Virus replication in tissues was measured by RT-qPCR at day 6 post-challenge only. Nasal cavity tissue RT-qPCR copy numbers were expressed as log_10_ values. For throat swab PCR data, the geometric mean of PCR copy numbers for days 1 to 6 post-challenge was taken for each animal and expressed as a log_10_ value, generating a single value measuring shedding over time for each animal. Both data sets were then tested using a one-way ANOVA followed by a comparison of study group means using the Tukey Honest Significant Difference test. All statistical analysis was carried out in JMP v14.3 (SAS Institute Inc.). Statistical significance are shown on figures by horizontal lines and P-value indicators: *P < 0.05, **P < 0.01, ***P < 0.001, ****P < 0.0001. Columns and error bars for all virus titer data show group geometric mean and interquartile range. To enable visualization of histopathology parameters from individual animals across the various study endpoints, all data were expressed as heatmaps.

## Acknowledgments

We thank Sarah Woods for FFA titration of vaccine formulation. We thank Laura Hunter and Chelsea Kennard for providing Histology support and expertise. Thank you to Marilyn Aram, Alastair Handley, Daniel Knott, Breeze Cavell, Stephen Thomas, Oliver Skinner and Thomas Bean for assistance with sample processing and all Biological Investigations Group staff for animal husbandry and assistance with *in vivo* procedures.

## Conflicts of interests

At the time of data generation for this publication, Katarzyna E. Schewe, Jonathan Crowe and Oliver Dibben were employees and shareholders of AstraZeneca plc.

AstraZeneca plc are the manufacturers of Fluenz® Tetra/FluMist® Quadrivalent intranasal influenza live virus vaccine.

## Contributions

Conceptualization: O.D., A.C.M., M.W.C., C.J.W.

Methodology: K.A.R., K.E.S.

Investigation: K.A.R., J.P., R.W., J.F.S., E.R., S.F., R.H.

Formal Analysis: K.A.R., K.E.S., J.C., J.P., R.W., J.F.S., E.R.

Writing – Original Draft: K.A.R., K.E.S.

Visualization: K.A.R., K.E.S.

Supervision: K.A.R., K.E.S., O.D.

Project Administration: O.D., K.E.S., K.A.R., Y.H., C.J.W.

Funding Acquisition: O.D.

Review & Editing: all authors

