## Supplementary Material for "Sequential delivery of LAIV and SARS-CoV-2 in the ferret model can reduce SARS-CoV-2 shedding and does not result in enhanced lung pathology"

### Supplementary Figures and Tables

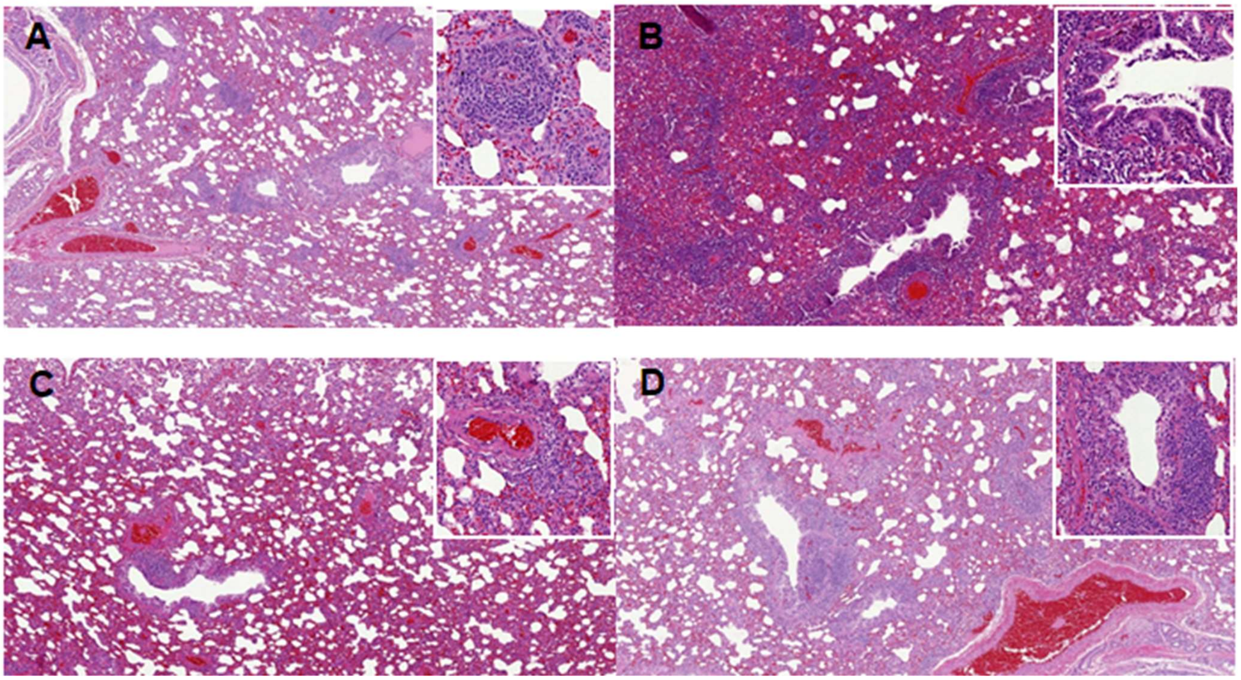


**Supplementary Figure 1. Histopathological lung changes.** A) SARS-CoV-2/LAIV, B) LAIV/SARS-CoV-2, C) Mock/SARS-CoV-2 and D) LAIV/Mock. Microscopic inflammatory changes in the parenchyma and airways. Insets show perivascular cuffing (A and C) and bronchiolar and peribronchiolar inflammatory infiltration (B and D). HE. 200 x magnification, inset 600 x magnification.

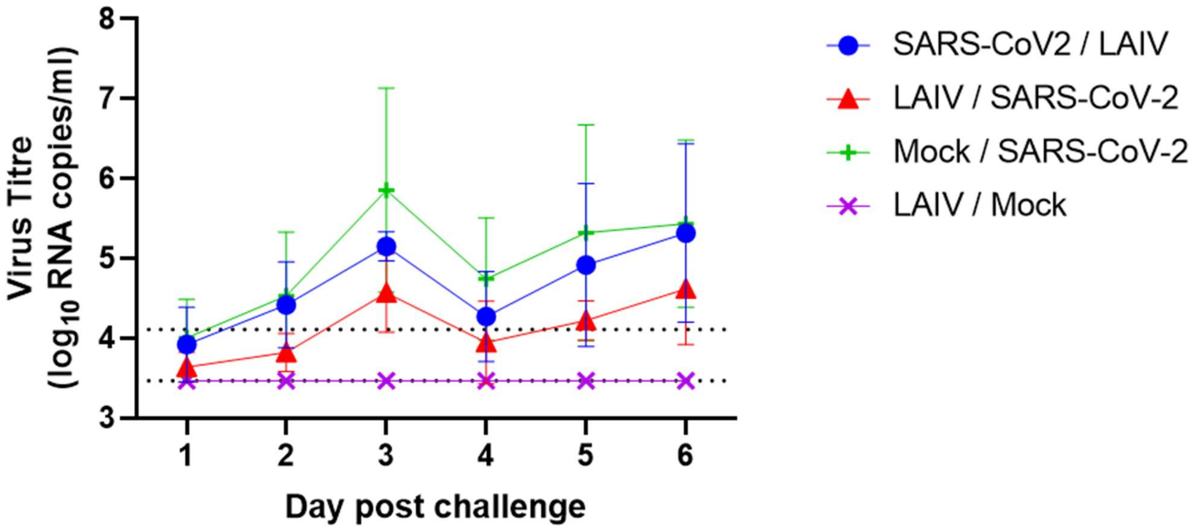


**Supplementary Figure 2. SARS-CoV-2 shedding over six days.** The throat swabs were taken at days 1, 2, 3, 4, 5 and 6 post challenge and SARS-CoV-2 viral loads were determined by RT-qPCR. The symbols show the viral RNA levels as the median of six ferrets with standard deviation. Horizontal dotted lines show the lower limit of quantification (LoQ=4.11 $\log_{10}$ RNA copies/ml/day) and the lower limit of detection (LoD=3.47 $\log_{10}$ RNA copies/ml/day).

**Supplementary Table 1. Summary of average lesion severity and viral RNA scoring in the lung and nasal cavity.**

| Group | Animal ID | Lung (left/right lobes) | | | | | | Nasal cavity | |
|---|---|---|---|---|---|---|---|---|---|
| | | Inflammation of alveolar walls/spaces (Focal alveolitis)* | Perivascular lymphocytic cuffing* | Bronchiolar inflammation (Bronchiolitis)* | Bronchial inflammation (Bronchitis)* | Viral staining intensity (SARS-CoV-2) | Total lung score** | Epithelial inflammation/ degeneration | Viral staining intensity (SARS-CoV-2) |
| SARS-CoV-2/LAIV | 8561 | 0.8 / 0.9 | 0.5 / 0.9 | 0.3 / 0.3 | 0.0 / 0.3 | 0 | 4 | 2.5 | 3 |
| | 7853 | 0.8 / 1.3 | 1.3 / 1.9 | 1.0 / 1.8 | 1.0 / 1.3 | 0 | 10.4 | 4 | 1 |
| | 11678 | 1.8 / 1.1 | 2.0 / 1.6 | 2.0 / 1.4 | 1.3 / 0.1 | 0 | 11.3 | 3.5 | 2 |
| | 7754 | 0.8 / 0.4 | 1.0 / 0.9 | 0.0 / 0.4 | 0.0 / 0.0 | 0 | 3.5 | 2.5 | 1 |
| | 10721 | 1.3 / 1.3 | 1.8 / 2.0 | 1.5 / 2.0 | 0.5/ 0.8 | 0 | 11.2 | 2 | 2.5 |
| | 8403 | 1.5 / 1.8 | 1.0 / 1.0 | 0.8 / 0.3 | 0.0 / 0.0 | 0 | 6.4 | 3.5 | 0 |
| LAIV/SARS-CoV-2 | 8656 | 1.8 / 1.4 | 1.8 / 2.5 | 1.5 / 2.3 | 1.0 / 0.6 | 0 | 13 | 3 | 0 |
| | 8666 | 1.8 / 1.5 | 3.5 / 2.3 | 2.8 / 1.9 | 1.3 / 0.9 | 0 | 16 | 4 | 0 |
| | 87853 | 1.3 / 1.0 | 1.5 / 1.0 | 1.5 / 1.0 | 1.0 / 1.0 | 0 | 9.3 | 3.5 | 0 |
| | 8433 | 1.3 / 1.3 | 1.8 / 1.0 | 2.0 / 1.3 | 0.8 / 0.9 | 0 | 10.4 | 4 | 0 |
| | 8616 | 0.8 / 0.6 | 1.3 / 1.8 | 0.8 / 1.5 | 1.0 / 0.8 | 0 | 8.6 | 3.5 | 0 |
| | 8676 | 1. 0 / 0.9 | 2.0 / 2.4 | 1.8 / 2.6 | 0.8 / 1.0 | 0 | 12.5 | 4 | 0 |
| Mock/SARS-CoV-2 | 8019 | 0.8 / 0.9 | 1.3 / 1.1 | 0.5 / 0.9 | 0.0 / 0.0 | 0 | 5.5 | 1.5 | 0 |
| | 8351 | 1.3 / 1.3 | 0.8 / 1.8 | 2.0 / 1.6 | 1.0 / 0.8 | 0 | 10.6 | 2.5 | 2.5 |
| | 8660 | 1.3 / 0.6 | 1.8 / 0.8 | 0.8 / 0.1 | 0.8 / 0.6 | 0 | 6.8 | 2 | 2.5 |
| | 11283 | 0.8 / 1.0 | 1.0 / 1.1 | 1.8 / 1.3 | 1.3 / 1.0 | 0 | 9.3 | 3 | 3 |
| | 8164 | 0.3 / 0.8 | 0.5 / 1.3 | 0.8 / 0.9 | 0.3 / 0.1 | 0 | 5 | 1 | 0 |
| | 7900 | 1.0 / 0.8 | 1.4 / 1.4 | 1.5 / 1.4 | 1.0 / 0.9 | 0 | 9.4 | 1 | 0.5 |
| LAIV/Mock | 7925 | 1.3 / 2.0 | 3.0 / 2.9 | 2.5 / 2.5 | 1.0 / 0.8 | 0 | 16 | 1 | 0 |
| | 8407 | 1.3 / 1.1 | 1.3 / 1.9 | 2.3 / 1.8 | 0.8 / 0.9 | 0 | 11.4 | 1 | 0 |
| | 8172 | 0.5 / 1.6 | 1.0 / 2.9 | 0.3 / 1.8 | 0.3 / 0.5 | 0 | 8.9 | 0.5 | 0 |
| | 10392 | 1.0 / 1.3 | 1.8 / 2.6 | 2.3 / 2.5 | 0.8 / 1.0 | 0 | 13.3 | 1 | 0 |
| | 7939 | 1.5 / 1.5 | 2.3 / 2.8 | 2.3 / 2.6 | 0.8 / 1.0 | 0 | 14.8 | 2 | 0 |
| | 8119 | 0.5 / 1.0 | 1.3 / 0.8 | 1.3 / 0.6 | 0.5 / 0.4 | 0 | 6.4 | 2.5 | 0 |

**Key:** *Mean scores for left / right lung lobes; ** sum of mean scores for left / right lung lobes

**Supplementary Table 2. Scoring criteria for the subjective assessment of microscopic changes in lung and nasal cavity of ferrets infected with SARS-CoV-2.**

| Location | Lesion | Score 0 (normal) | Score 1 (minimal) | Score 2 (mild) | Score 3 (moderate) | Score 4 (marked) |
|---|---|---|---|---|---|---|
| Lung | Infiltration of alveolar walls and spaces by inflammatory cells (focal alveolitis) | None | Rare foci | Increased numbers, up to 25% of the slide affected | Frequent areas between 26-50% of the slide affected | Numerous areas; over 50% of the slide affected |
| | Perivascular inflammatory infiltrates (cuffing) | None | Occasional, up to 5% of vessels | Increased numbers, up to 25% of vessels | Numerous cuffs; between 26-50% of vessels | Over 50% of all vessels |
| | Bronchial inflammation with presence of exudates and/or inflammatory cell infiltration | None | Occasional, (1 or 2) bronchi affected | Present in multiple airways; up to 25% of bronchi affected | Present in multiple airways; between 26-50% of bronchi affected | Present in multiple airways; over 50% of bronchi affected |
| | Bronchiolar inflammation with presence of exudates and/or inflammatory cell infiltration | None | Occasional, (1 or 2) bronchioles affected | Present in multiple airways; up to 25% of bronchioles affected | Present in multiple airways; between 26-50% of bronchioles affected | Present in multiple airways; over 50% of bronchioles affected |
| Nasal cavity | Inflammatory cell infiltration +/- epithelial degeneration/necrosis | None | Rare inflammatory cell infiltrates +/- epithelial degeneration/necrosis up to 5% of slide affected | Multifocal inflammatory cell infiltrates +/- epithelial degeneration/necrosis, between 6-25% of the slide affected | Multifocal inflammatory cell infiltrates +/- epithelial degeneration/necrosis between 26-50% of the slide affected | Multifocal inflammatory cell infiltrates +/- epithelial degeneration/necrosis; over 50% of the slide affected |
